# Home-cage Monitoring Ascertains Signatures of Ictal and Interictal Behavior In Mouse Models of Generalized Seizures

**DOI:** 10.1101/740407

**Authors:** Miranda J. Jankovic, Paarth P. Kapadia, Vaishnav Krishnan

**Affiliations:** Department of Neurology, Laboratory of Epilepsy and Emotional Behavior Baylor Comprehensive Epilepsy Center, Baylor College of Medicine One Baylor Plaza MS NB302 Houston, TX 77030

**Keywords:** Epilepsy, pentylenetetrazole, interictal, home-cage monitoring, psychiatric comorbidities of epilepsy, ethologically valid, ethological approach

## Abstract

Epilepsy is a significant contributor to worldwide disability. In epilepsy, disability has two components: ictal (pertaining to the burden of unpredictable seizures and associated medical complications including death) and interictal (pertaining to more pervasive debilitating changes in cognitive and emotional behavior). In this study, we objectively and noninvasively appraise correlates of ictal and interictal disability in mice using instrumented home-cage chambers designed to assay kinematic and appetitive behavioral measures. We discover that in C57BL/6J mice, intraperitoneal injections of the chemoconvulsant pentylenetetrazole (PTZ) acutely result in complex and dynamic changes in movement and sheltering behavior that evolve (or kindle) with repeated daily injections, and which are separate from the occurrence of convulsions. By closely studying “interictal” periods (between PTZ injections), we identify a syndrome of nocturnal hypoactivity and increased sheltering behavior. We observe elements of this interictal behavioral syndrome in seizure-prone DBA/2J mice and in mice with a pathogenic *Scn1a* mutation (modeling Dravet syndrome). Through analyzing their responses to PTZ, we illustrate how convulsive severity and “behavioral” severity are distinct and independent aspects of overall seizure severity. Our results illustrate the utility of an ethologically centered automated approach to quantitatively appraise murine expressions of disability in mouse models of seizures and epilepsy. In doing so, this study highlights the very unique psychopharmacological profile of PTZ.

**Significance Statement:** Epilepsy is a brain disorder characterized by a pervasively increased risk to develop epileptic seizures. Sadly, patients with epilepsy also experience high rates of anxiety, depression and other psychiatric symptoms that significantly increase overall disability. While many mouse models of seizures and epilepsy exist, we need improved techniques to measure how new treatments impact not only seizure occurrence, but also emotional changes that persist in between seizures. In this study, we apply the technique of home-cage monitoring to clarify precisely how spontaneous mouse behavior is altered in three distinct epilepsy models. Our work illustrates the importance of an ethologically centered appreciation of neuropsychiatric disability in mice and clarifies a new approach to the measurement of “seizure severity”.

## Introduction

Epilepsy, or the epilepsies, are an etiologically diverse group of disorders defined by an enduring predisposition to develop epileptic seizures, and by associated neurobiologic, cognitive/psychological and social consequences (1). Epilepsy syndromes contribute significantly to worldwide disability and premature mortality (2, 3). In cross-sectional studies of adults with epilepsy, self-reported assessments of disability show some correlation to seizure frequency/recency, but also correlate with anticonvulsant side effects and the intensity of comorbid mental health impairments, including anxiety, depression and/or sleep disturbances (4, 5). These comorbidities magnify stigma, complicate anticonvulsant selection (6) and are associated with lower rates of seizure freedom following epilepsy surgery (7). Understanding the etiology of these comorbidities and their complex relationships with the underlying epileptogenic lesion/network remains an important consensus research benchmark (8), and is essential for intelligently designing novel therapeutics that comprehensively address ictal and interictal contributors to disability.

With the increasing appreciation of epilepsy as a spectrum disorder (9), rodent studies employing genetic and/or pharmacological manipulations popularly report seizure-related measures as well as measures of behavioral comorbidity. The gold standard for the detection of spontaneous seizures is video electroencephalography (vEEG), obtained via surgically implanted tethered or wireless EEG electrodes. When spontaneous seizures are extremely rare or absent (10–12), conclusions about “enhanced seizure severity” or “lowered seizure thresholds” are inferred from responses to experimentally induced acute seizures. One such widely popular induction agent is pentylenetetrazole (PTZ), a gamma-aminobutyric acid type A (GABA-A) receptor antagonist (13), used briefly in the past to induce habitual seizures in humans with rare events (14). In mice, PTZ injections produce a spectrum of behavioral changes, ranging from hypoactivity/immobility, clonic or myoclonic seizures, to tonic-clonic convulsions (15, 16). These seizures recapitulate the semiology, electroencephalographic characteristics (17) and pharmacological responsiveness (18) of seizures with a generalized onset. Differences in seizure severity are deduced from (i) subjectively scored ordinal scales of convulsive severity (e.g., Racine scale) that are inconsistently applied (19–22), (ii) measures of convulsion latency or duration (17, 18), and/or (iii) rates of frank seizure-induced mortality (11). These measures are based on the assumption that more severe seizures in mice are those that are convulsive and/or lethal, arise with a shorter latency and/or are of longer duration. However, they provide no insights into how seizures interrupt/disrupt a mouse’s interaction with its environment. They also offer no information about the duration or intensity of post-ictal impairment, quantities that are also critical contributors to epilepsy severity and which may also possess translational value.

Aside from measurements of learning and memory, assessments of interictal emotional behavior in rodents have relied heavily on phenotyping batteries (23–25) to identify features of “depression-related” or “anxiety-related” behavior extrapolated from assessments of behavioral despair (e.g., forced swim test) or exploration (e.g., open field / elevated plus maze tests). While these tests are certainly convenient and offer a reasonably high throughput, they are short in duration (5-10 minutes long) and are often measured during the biological night (when lights are on). Virtually all of these tasks necessitate human presence/interference, which may directly impact reproducibility through inconsistent olfactory cues, variations in technique and bias (26, 27).

In this study, we utilize an established home-cage monitoring platform (28, 29) to obtain objective and unbiased continuous measures of ictal and spontaneous interictal murine behavior with negligible human presence in three distinct mouse models. With PTZ injections, we demonstrate how acute convulsive severity is independent and distinct from “behavioral” severity. Further, we extract patterns of interictal disability from prolonged (≳19h) recordings of home-cage behavior that emphasize the nocturnal/active phase.

## Results

### Spontaneous Home-cage Behavior in Wildtype C57BL6/J Mice

Adult C57BL/6J mice were individually housed in home-cage chambers (30×30×47cm) containing an infrared-lucent shelter, two lickometered water spouts (water Vs 0.8% sucrose) and a beam-break metered food hopper. Before intraperitoneal injections, we obtained two days of baseline recordings (lights are off between 1700-0500, defined as the *“*active phase”). Here, we profile data from the second day (Fig. 1A). Between 1500-1200, mice accumulated an average total distance of ∼850m (Fig. 1B). Heat maps of position probability during 6h-long epochs demonstrated peaks at the food hopper and shelter that inversely varied in prominence over the day. Averaged time budgets (30) quantified this effect (Fig. 1C): these were calculated by summarizing total durations (per epoch of time) spent at the feeder, water spouts, shelter or engaged in “other” behaviors (time spent *not* feeding, drinking or sheltering). Consumptive and sheltering behavior were generally synchronized with locomotor activity (Fig. 1D, E), and mice displayed an average sucrose preference of ∼80%. To measure “sleep” noninvasively, we identified contiguous periods of lack of movement that were ≥ 40s in duration (see Fig. S1A for a pharmacological validation of this technique with diazepam). This approach displays ∼90% agreement with sleep states derived from electroencephalography-electromyography but does not discriminate rapid eye movement (REM) sleep from non-REM sleep (31, 32). C57BL/6J mice on average spent ∼10.3h in “sleep”, with sleep bouts (mean duration ∼263s, Fig. 1F) occurring throughout the day consistent with prior reports (33–35). Together, these data recapitulate (in a laboratory setting) natural circadian patterns of foraging and sheltering behaviors of mice in the wild, achieved entirely through automation and without concurrent human presence or invasively placed telemetry implants (36).

**Fig. 1.**
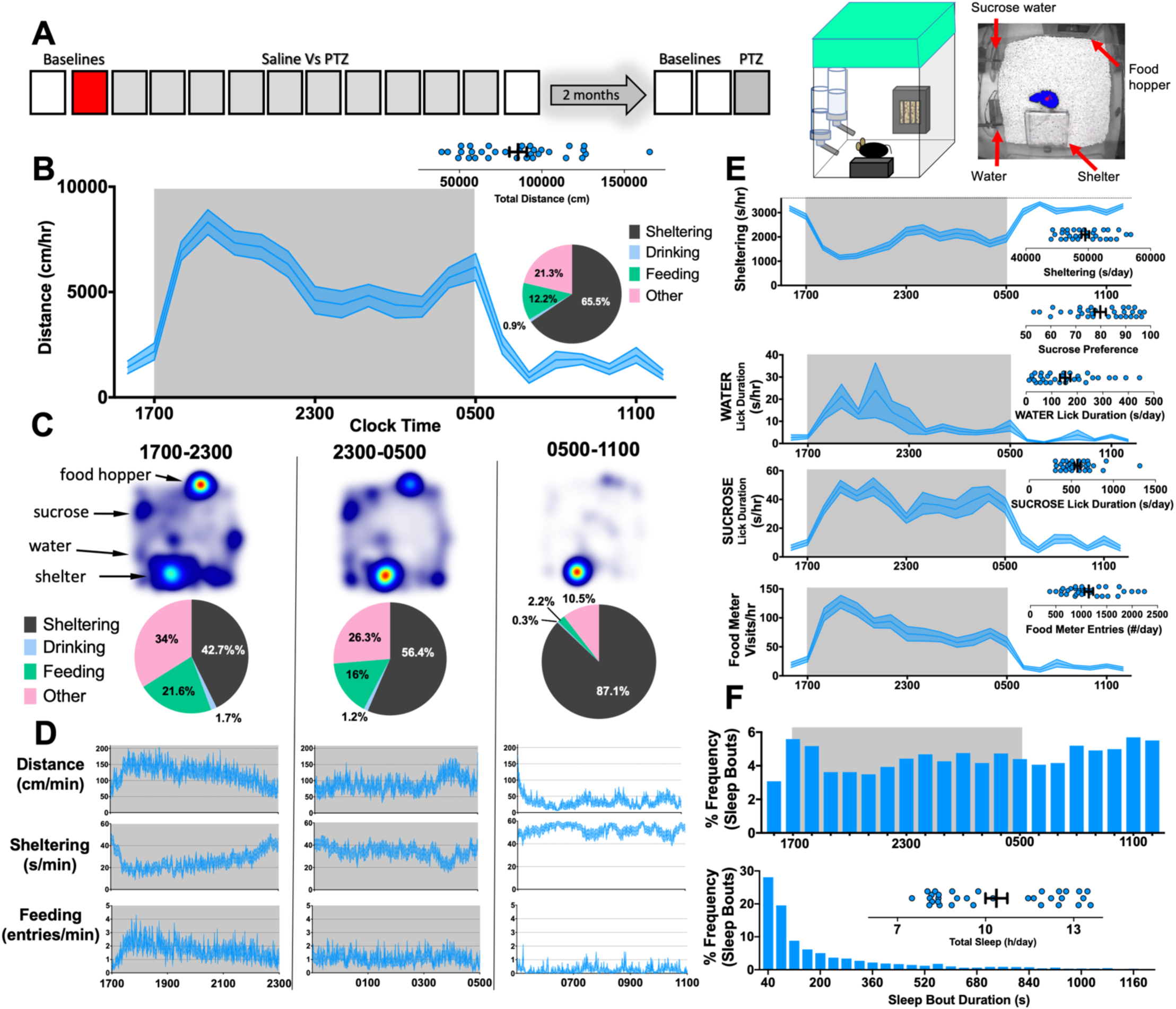
Home-cage behavior in C57BL/6J mice. **(A)** Experimental protocol for 8-week old C57BL/6J mice (n=32, 16 female), with the epoch depicted by this figure in marked red. RIGHT: Cartoon showing home-cage configuration with a screen capture from an aerial infrared camera showing mouse body contour (blue) and centerpoint (red). **(B)** Distances moved (per hour) on baseline day 2, with shaded box depicting the active phase. INSET: time budget across this 21h recording period. **(C-D)** Heat maps, time budgets and behavioral quantities depicted over 6h long epochs where “other” is defined as time spent *not* sheltering, drinking or feeding. **(E)** Various parameters measured simultaneously with individual total values plotted in inset. **(F)** Percent frequency of sleep bouts as a function of time of day (TOP) and by duration of sleep bout (BOTTOM), with individual values obtained for total sleep (INSET). Mean ± standard of the mean (SEM) shown.

### Acute Effects of Intraperitoneal Pentylenetetrazole Injections

We randomized these mice into two sex-matched groups (Fig. S1B-E) to receive 10 daily intraperitoneal injections of either saline or PTZ at a subconvulsant dose of 30mg/kg (16, 19, 20). Injections were administered at ∼1200 for three reasons: (i) to respect how many epilepsy mouse models display peaks of spontaneous seizure occurrence during the inactive phase (37–39), (ii) to avoid light-induced circadian disruptions that invariably occur when injections must be administered with lights off, and (iii) to elaborately emphasize how daily seizures impact the entirety of active phase behavior. Immediately following injections, we closely analyzed 3h-long “ictal” recordings as well as more prolonged “interictal” recordings (1600 to 1100, Fig. 2A). On the first injection day (Fig. 2B), saline-treated mice displayed a gradual reduction in locomotor activity over ∼30 minutes, following which they retreated to a state of marked immobility (30-90 minutes) within shelters. Approximately 90 minutes following saline injection, mice resumed normal rates of daytime movement and sheltering behavior (compare with Fig. 1C). Over subsequent days, behavioral responses to saline injections remained fairly stereotyped (Fig. 2B-D, S2).

**Fig. 2.**
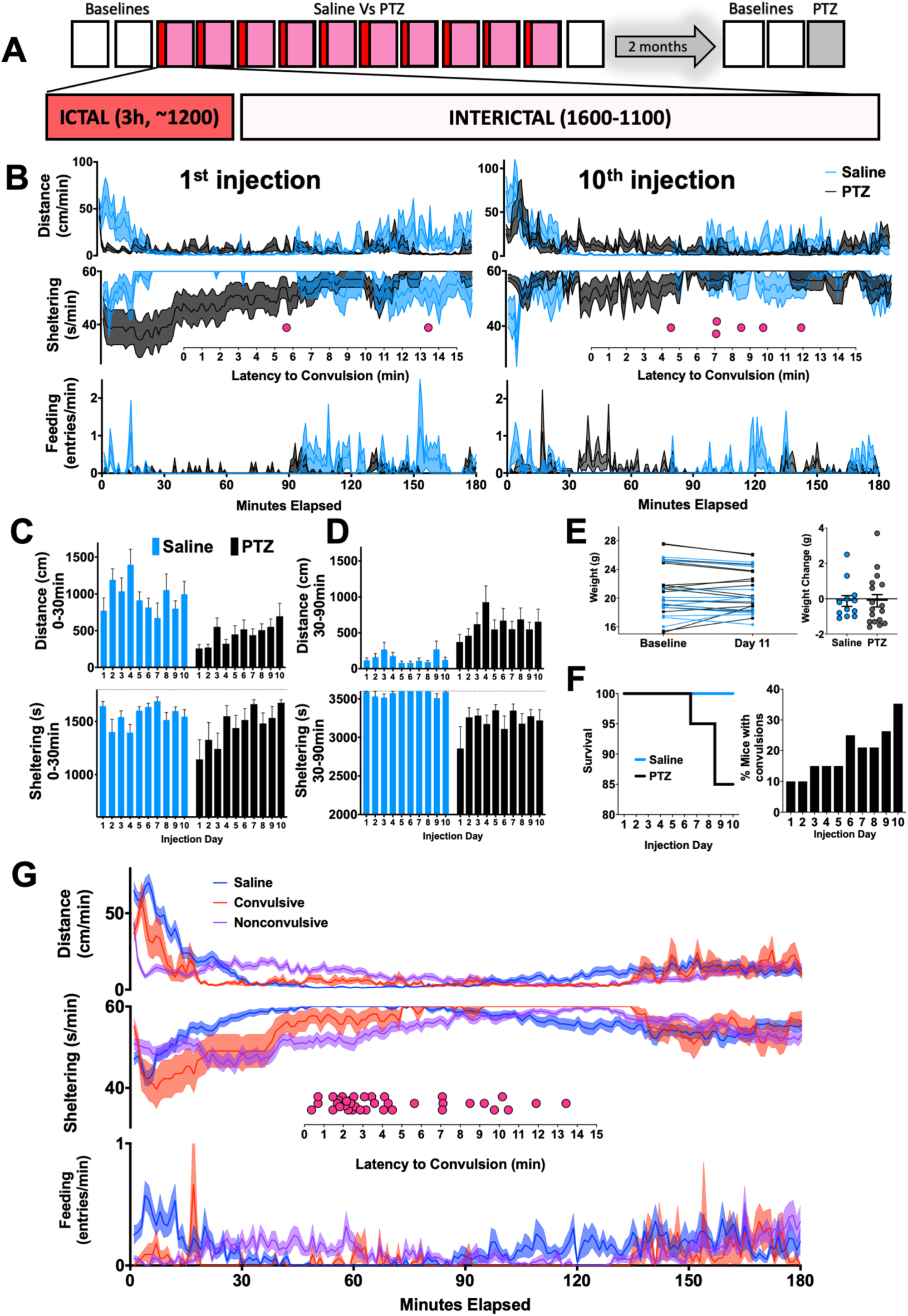
Acute changes in home-cage behavior. **(A)** 10 daily 3h-long “ictal” recordings beginning at approximately noon following injections of either saline (n=12) or 30mg/kg PTZ (pentylenetetrazole, n=20) **(B)** Changes in distance moved, sheltering and feeding observed during the 1^st^ and 10^th^ injections (see also Fig. S2). Dot plots depict latencies to convulsions. **(C)** In the first 30 minutes, total distance and sheltering time increased over the 10-day injection period (distance: group x day, F_9,261_ = 2.95, p<0.01, and sheltering: group x day, F_9,261_ = 2.00, p<0.05). **(D)** In the subsequent hour (30-90mins), total distances (group x day, F_9,261_ = 0.91, p>0.05) and sheltering (group x day, F_9,261_ = 0.95, p>0.05) did not significantly evolve. **(E)** Weight change between groups (p>0.5). **(F)** Survival and occurrence of convulsions over the 10-day protocol. **(G)** Ethograms averaged across all 10 days for saline injections (n=120), convulsive (n=33) and nonconvulsive (n=158) ictal recordings (inset: latency to convulsion across all days). Mean ± SEM shown.

In contrast, the first PTZ injection produced an early steep decline in locomotion associated with a significant sheltering deficit (Fig. 2B, Movie S1). With repeated PTZ injections, this early sheltering deficit and immobility response gradually improved (Fig. 2C). Between 30-90 minutes after the first injection, PTZ-treated mice displayed an unexpected relative increase in locomotion with continued deficits in sheltering (Fig. 2B). Relative hyperactivity and sheltering deficits (during this epoch) did *not* substantially evolve with repeated PTZ injections (Fig. 2D). Weight change, assessed over the 10 day protocol, was no different between these groups (Fig. 2E).

Through an integrated visualization of video and real-time measures of mobility (see Materials and Methods, Movie S2), we retrospectively identified the occurrence of convulsive seizures (37 in total), which as expected, increased in probability with repeated PTZ exposure (Fig. 2F, S2, Movie S3, S4). Convulsions occurred with a mean latency of 4.44 min, and three convulsions resulted in death. Mice that displayed any convulsions (9 out of 20) trended to be more hyperactive (at baseline) than those that never convulsed (Fig. S3A). 7 out of 20 mice displayed convulsions following the second or subsequent PTZ injections. When compared with the 11 mice that never convulsed, these mice displayed significantly lower amounts of sheltering between 30-90 minutes after their *first* injection of PTZ (Fig. S3B). Averaged across *all* 10 days (Fig. S3C), PTZ injections produced a fairly stereotyped deviation from saline injections. When this average response was separated by the presence or absence of a convulsion, distinct ethograms emerged, each uniquely different from that observed in saline-treated mice (Fig. 2G).

### The Interictal Behavioral Syndrome Induced by PTZ Injections

We obtained 19h long “interictal” recordings of spontaneous home-cage behavior after every injection to quantify more delayed behavioral differences (Fig. 3A). Saline injections alone reduced total daily distances to ∼600m/d, but this remained fairly stable with repeated injections. In contrast, PTZ-treated mice displayed a progressive relative decline in total locomotor activity (Fig. 3B) associated with increased total sheltering but without an increase in overall “sleep”, sucrose preference or total licking (Fig. S4A) On many but not all injection days, PTZ-treated mice displayed fewer entries into the food meter (Fig. 3B). Sucrose preference and total licking was similar in PTZ and saline groups (Fig. S4A). The 10^th^ interictal period is profiled in Fig. 3C, where prominent differences in home-cage behavior occurred during the active phase. PTZ-treated mice spent more time sheltering at the “cost” of reduced feeding and “other” behaviors (Fig. 3D), reflecting a more general reduction in home-cage exploration.

**Fig. 3.**
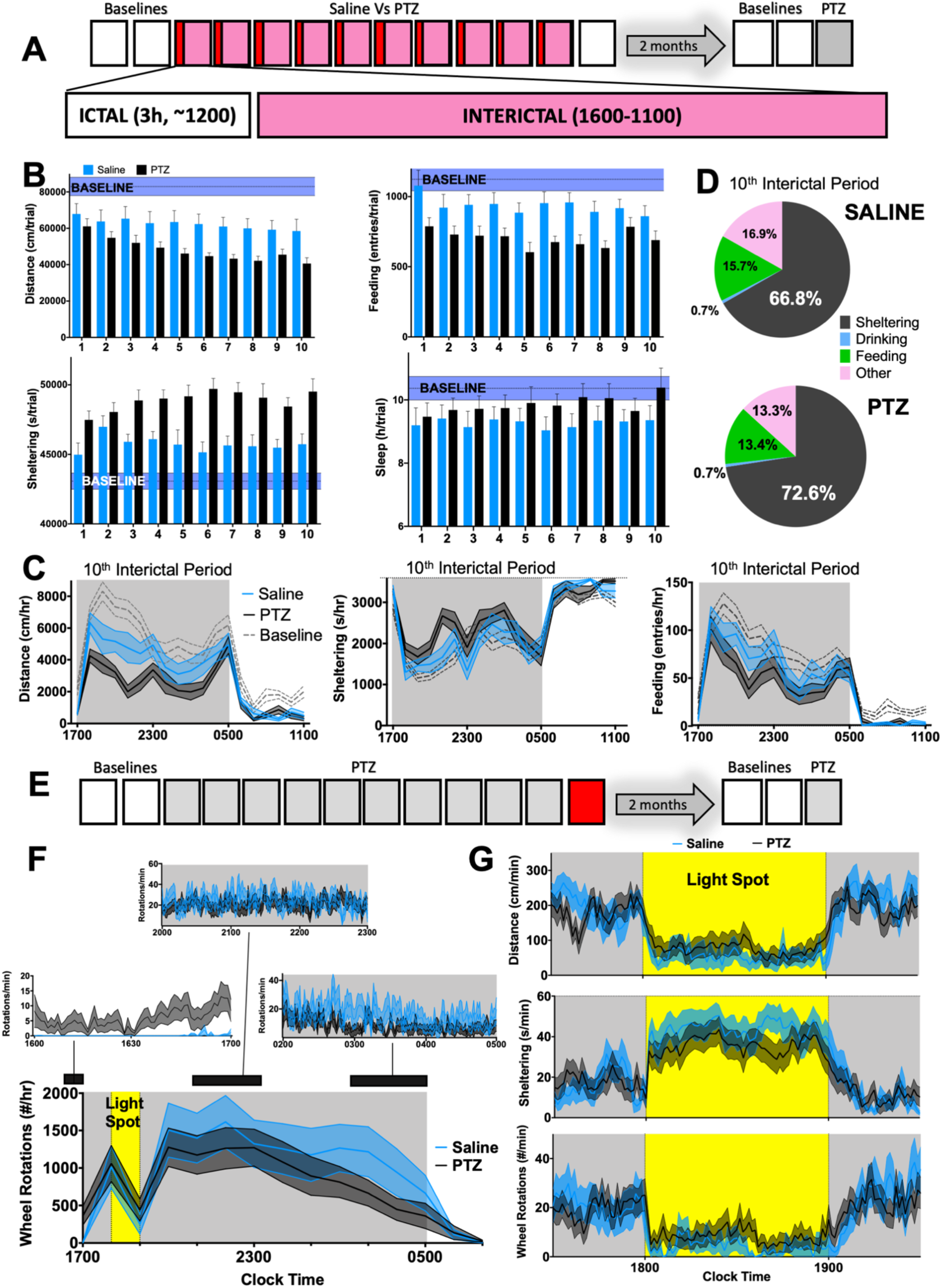
Daily PTZ injections produce an interictal syndrome of hypoactivity and increased sheltering. **(A)** 10 daily 16h-long “interictal” recordings beginning at 1600 (saline n=12, PTZ n=17-20) **(B)** PTZ injections produced hypoactivity (group x day, F_9,261_ = 1.64, p=0.1), increased sheltering (group x day, F_9,261_=2.17, p<0.05), without changing feeding (group x day, F_9,261_=0.63, p>0.5) or sleep (group x day, F_9,261_=0.66, p>0.5). Baseline ranges (corresponding to total measurements between 1600-1100 for day 2) are depicted in purple (mean ± SEM) **(C)** 10^th^ interictal period: PTZ resulted in hypoactivity (group x hour, F_18,486_=2.2, p<0.01), a trend for increased sheltering (group x hour, F_18,513_=1.47, p = 0.09) without altering feeding (group x hour, F_18,486_=0.97, p=0.4). **(D)** Averaged time budgets for C. **(E)** On the day after their last PTZ or saline injection, **(F)** voluntary wheel running was measured. PTZ-treated mice displayed increased wheel running during the first hour (saline 0.1±0.05 Vs PTZ 6.4±2 rotations/min, p<0.05), but similar wheel running during the mid-active phase (2000-2300, saline 24.1±5.3 Vs PTZ 20.6±4.2 rotations/min, p>0.1) or during the late active phase (0200-0500, saline 15.7±4.7 Vs PTZ 8.1±1.8 rotations/min, p>0.5). **(G)** A light spot test produced similar changes in activity and sheltering in PTZ and saline-treated mice. Mean ± SEM shown.

We incorporated two additional perturbations the following day (Fig. 3E). First, we introduced running wheels just before 1600. During the first hour of wheel exposure, PTZ-treated mice accumulated significantly more wheel rotations (Fig. 3F). At active phase onset, both groups mounted a robust increase in wheel running (∼1000 rotations/hr or ∼471m/hr, Fig. 3G). At 1800, we applied a 60-minute long “light spot” stimulus (40), which interjects a conflict between nocturnal foraging behavior and light aversion. This light stimulus attenuated locomotion and enhanced shelter entry, but saline and PTZ-treated mice displayed largely similar responses (Fig. 3H).

### Delayed Effects on Seizure Severity and Anticonvulsant Pre-Treatment

The interictal syndrome identified in the midst of repeated PTZ injections was not observed when the same mice were studied two months later (Fig. 4A-C, Fig. S4B). After these baseline recordings, all mice received a final injection of PTZ (30mg/kg). Previously saline-treated mice displayed marked immobility. In contrast, previously PTZ-treated mice displayed greater convulsive severity and early relative hyperactivity (Fig. 4D, E), similar to the “kindled” response seen in Fig. 2B. Next, in order to appreciate whether anticonvulsant pre-treatment would impact home-cage responses to PTZ, a separate group of 8-10week old mice were acclimated to home-cages for 48h. Then, mice received either an injection of saline or sodium valproate [200mg/kg] (18, 41), followed 1h later by a 30mg/kg PTZ injection. Valproate pre-treatment did not significantly affect behavioral or convulsive indices of seizure severity (Fig. 4F). The following day, we applied similar saline or valproate injections an hour before a convulsant dose of PTZ (60mg/kg). In this trial, valproate resulted in significantly fewer convulsions, but dynamic measures of distance and sheltering behavior were similar between groups.

**Fig. 4.**
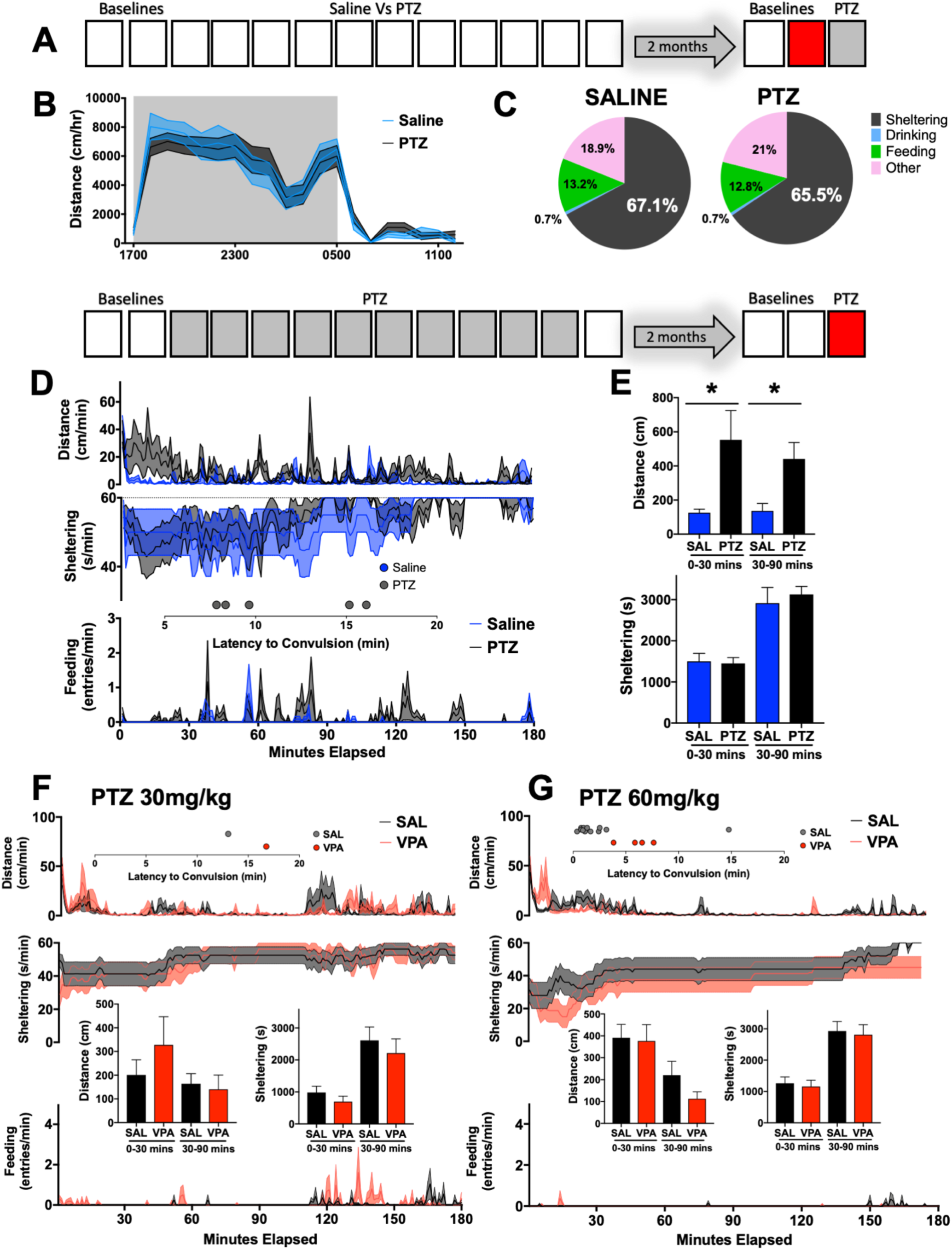
Home-cage behavior after seizure remission and anticonvulsant treatment. **(A)** Two months after the cessation of daily injections, mice were reintroduced into home-cage monitoring chambers. **(B)** Overall distances moved between previously saline (n=12) and PTZ-treated (n=17) mice were no different (group x hour, F_19,513_= 0.44, p>0.5. **(C)** Time budgets for epoch shown in B. **(D)** Final injection of PTZ (30mg/kg): previously saline-treated mice displayed marked immobility, whereas previously PTZ-treated mice displayed relative hyperactivity together with convulsions (PTZ 5/17 vs Saline 0/12, χ^2^ = 4.2, p<0.05). **(E)** Quantification of findings in D. Distances, sheltering and feeding responses following a **(F)** 30mg/kg or **(G)** 60mg/kg PTZ injection approximately one hour after an intraperitoneal injection of saline or 200mg/kg VPA (sodium valproate, n=16 [8 females]/group). Valproate-treated mice displayed significantly fewer convulsions (4/16 Vs 13/16, χ^2^ = 10.1, p<0.01). * p<0.05. Mean ± SEM is shown.

### Comparing C57Bl/6J and DBA/2J mice

Compared with C57BL/6J mice, mice of the DBA/2J strain display greater seizure severity scores following exposure to PTZ and other GABAergic antagonists (15, 42). To explore how such differences in seizure threshold may associate with changes in home-cage behavior, we compared age and sex-matched C57BL/6J and DBA/2J mice bred under identical conditions (Fig. 5A). In addition to hypoactivity, DBA/2J mice also displayed lower entries but a greater duration of time at the food hopper (Fig. 5B-D). DBA/2J mice had shorter mean durations of “sleep” bouts without significant differences in overall “sleep” (Fig. 5E). In response to a light spot (Fig. 5F), DBA/2J mice displayed rapid and sustained shelter entry together with prominent locomotor suppression for the duration of the light stimulus. Importantly, at this age, both strains are similar in light detection and visual acuity (43).

**Fig. 5.**
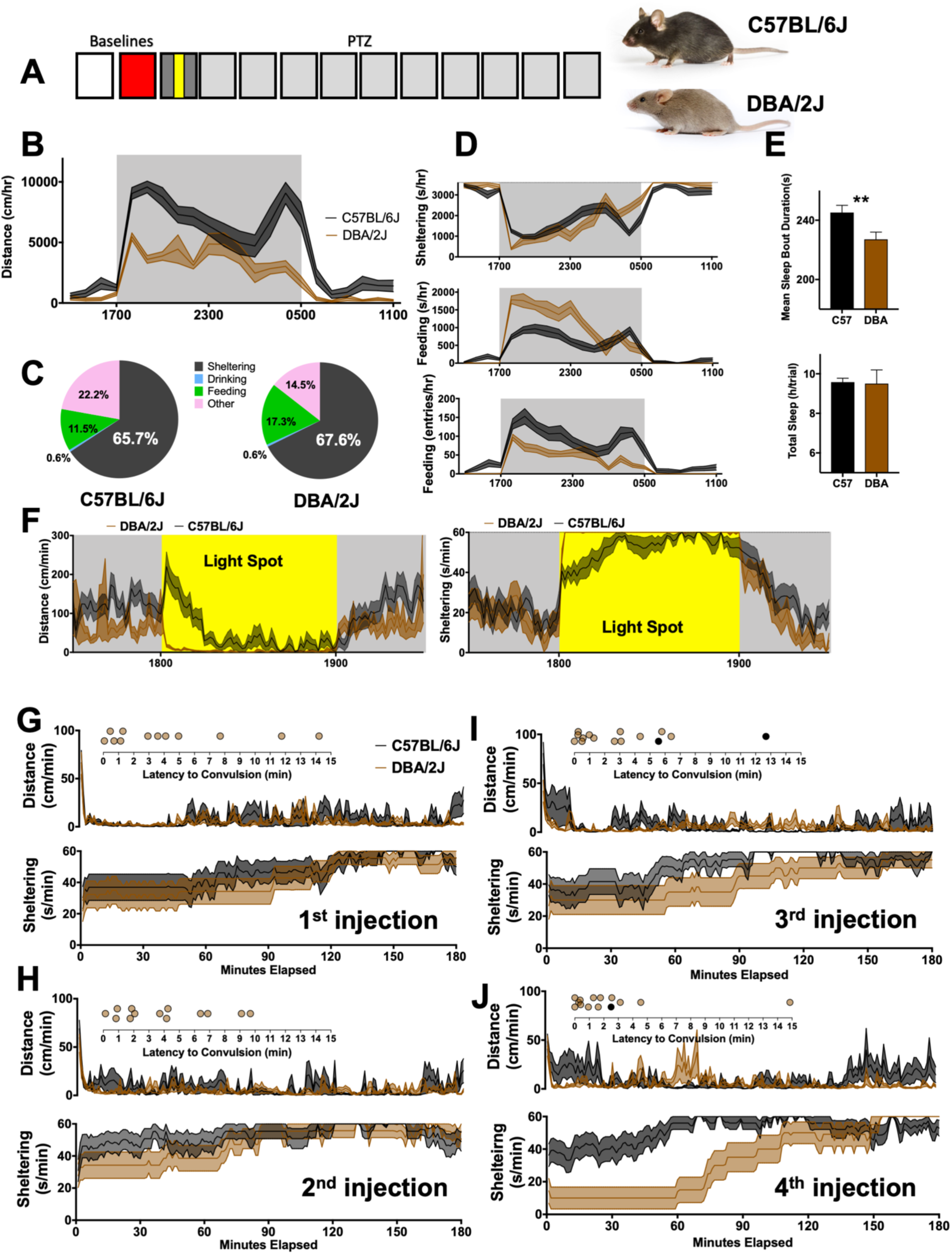
C57BL/6J Vs DBA/2J. **(A)** 8-10week old C57BL/6J (n=15, 7 female) and DBA/2J (n=15, 7 female) were directly compared. Photographs with permission from © The Jackson Laboratory. **(B)** On the second baseline day, DBA/2J mice were markedly hypoactive (genotype x time, F_21,588_=5.9, p<0.0001). Overall time budgets are shown in **(C)**. **(D)** DBA/2J mice also displayed significantly different patterns of sheltering (genotype x hour, F_21,588_=4.3, p<0.0001), increased feeding duration (genotype x hour, F_21,588_=8.7, p<0.0001) and decreased feeding entries (genotype x hour, F_21,588_=3.8, p<0.0001). **(E)** Mean sleep bout duration and total sleep time. **(F)** During the light spot hour, DBA/2J mice displayed rapid and committed shelter entry and immobility. **(G-I)** Distances and sheltering patterns for the first-through fourth daily injections of PTZ in ∼10week old female DBA/2J (n=14) and C57BL/6J (n=13) mice with insets depicting the occurrence of convulsive seizures (see also Fig. S5). ** p<0.01. Mean ± SEM is shown.

In preliminary experiments, a high rate of PTZ-induced mortality was seen in male DBA/2J mice: in a cohort of 8 male DBA/2J mice, only 2 survived past the sixth PTZ injection. Therefore, here we present data from only female mice (of both inbred strains). Following the first PTZ injection, C57BL/6J and DBA/2J mice displayed similar patterns of locomotor activity and sheltering behavior, but DBA/2J displayed a significantly higher incidence of convulsions (Fig. 5G). Over subsequent days of testing, DBA/2J continued to display the vast majority of convulsions and experienced seizure-induced mortality at a greater rate (Fig. 5H-J, Fig. S5). In several but not all of their subsequent PTZ injections, DBA/2J mice exhibited marked delays to shelter re-entry (Fig. S5).

### Ictal and Interictal Measures in a Mouse Model of Dravet Syndrome

Dravet syndrome (DS) is a neurodevelopmental disorder characterized by intellectual disability, autism spectrum disorder and epilepsy (44). Mutations in the voltage gated sodium channel subunit *SCN1A* are found in approximately 80% of patients with DS (45) and mice engineered to heterozygously express a clinically observed truncation mutation in *Scn1a* (R1407X, hereafter referred to as Scn1a^+/-^) display rare spontaneous seizures with a normal interictal electroencephalogram (46–48). Adult Scn1a^+/-^ mice have been reported to display altered anxiety-related behaviors, impaired sociability and spatial memory (49). In our home-cage chambers, compared with wildtype littermates, Scn1a^+/-^ mice displayed early active phase hypoactivity, without alterations in feeding and sheltering (Fig. 6A). Like PTZ-treated mice (Fig. 3D), Scn1a^+/-^ mice spent more time sheltering at the cost of reduced feeding (Fig. 6B). Overall licking was reduced without affecting sucrose preference, and Scn1a^+/-^ mice displayed significantly higher total “sleep” and longer bouts of “sleep” (Fig. 6C). In response to a light spot stimulus (40), Scn1a^+/-^ mice and WT littermates performed comparably (Fig. 6D). In contrast, Scn1a^+/-^ mice displayed hypoactivity and accelerated shelter entry following a temporary (2h long) “cage swap” (Fig. 6E). Finally, in response to 30mg/kg of PTZ, Scn1a^+/-^ mice displayed prominent immobility during both early (0-30min) and late (30-90min) epochs (Fig. 6F, G). Scn1a^+/-^ mice also displayed a prominent suppression of feeding over the entirety of the 3h long observation period. Only one Scn1a^+/-^ mouse convulsed.

**Fig. 6.**
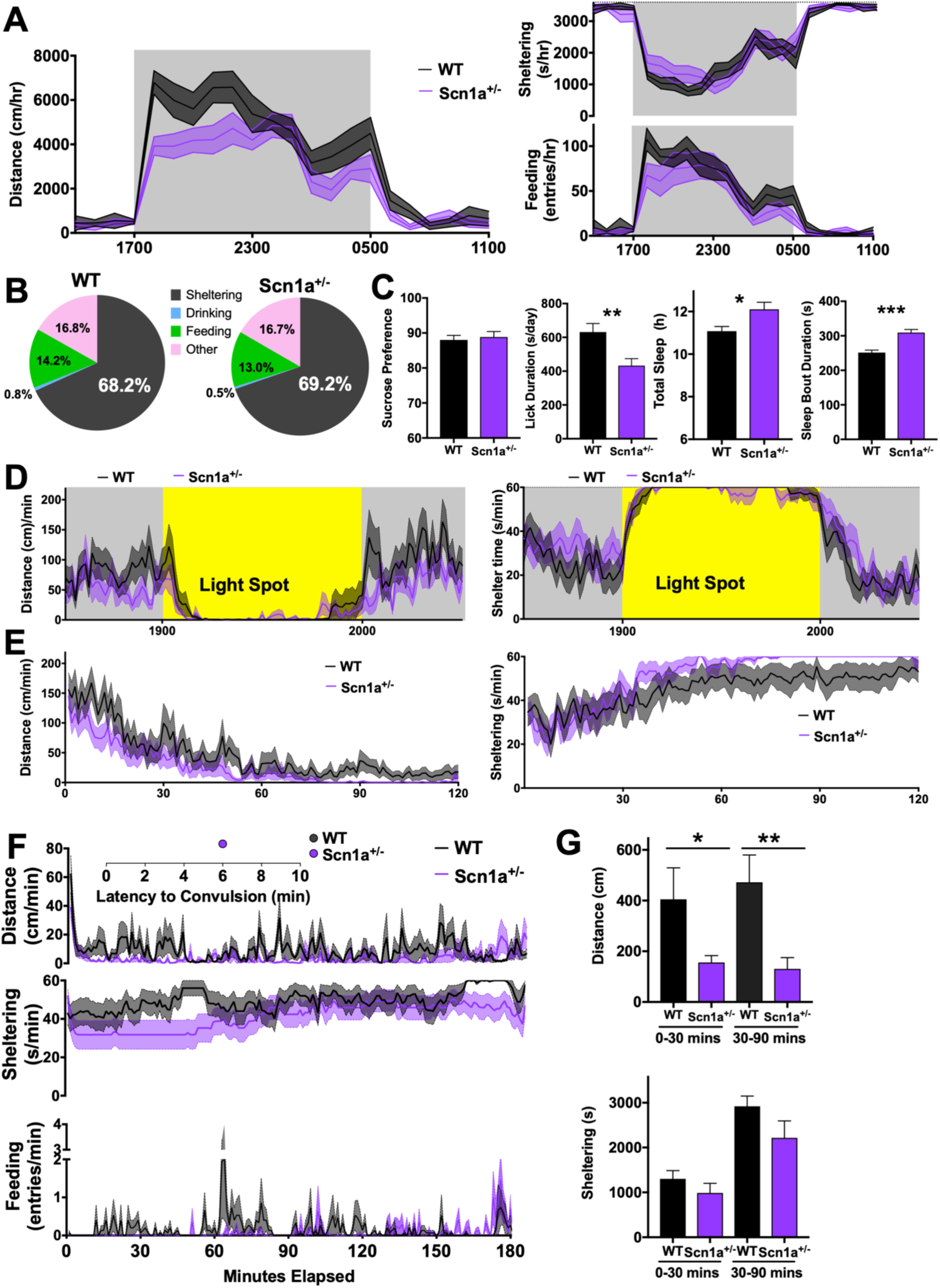
Home-cage behavior in Scn1a^+/−^ mice. **(A)** 8-10week-old Scn1a^+/−^ mice (n=15, 8 male) were hypoactive compared with littermate WT mice (n=13, 8 male, genotype x hour F_21,546_=2.21, p<0.01) without significant differences in sheltering or feeding entries (p>0.1). **(B)** Time budgets for A. **(C)** Scn1a^+/−^ mice displayed lower overall total licking without a change in sucrose preference, together with increased totalsleep time and mean sleep bout durations. **(D)** Light spot stimulation revealed no differences, but **(E)** with a day time “cage-swap”, Scn1a^+/−^ mice displayed more rapid locomotor habituation and accelerated shelter entry. **(F)** Following PTZ (30m/kg), Scn1a^+/−^ mice displayed greater immobility, quantified in **(G)**. WT=wildtype. *p <0.05, **p<0.01. Mean + SEM is shown.

## Discussion

In this study, we applied home-cage monitoring in mice to describe and appraise the acute severity of induced seizures as well as the severity of more pervasive changes in behavior that arise with recent seizures or with pervasively increased seizure risk (i.e., epilepsy). Home-cage monitoring technology has unearthed important behavioral phenotypes in mouse models of autism (50–52), neuromuscular and movement disorders (53, 54) and Alzheimer-related cognitive impairment (55, 56). The capacity to measure multiple variables simultaneously over prolonged recording periods avoids biased assumptions about *when* (i.e., the time of day) or *how* (i.e., which behavioral axis) a genetic or pharmacological manipulation may be symptomatic (26). Such home-cage assessments may provide superior translational value, especially when aligned with human data captured by wearable devices designed to quantify sleep, sedentariness and frailty. Since the behavioral expression of a static epileptogenic structural or genetic lesion may depend on the timing of the most recent seizure, we submit that long term home-cage monitoring (especially when combined with electroencephalography) is particularly well-suited to study how seizure occurrence impacts interictal behavior, or vice versa.

### Behavioral Severity is Independent of Convulsive Severity

The first objective was to apply home-cage monitoring to study seizure severity, an endophenotype central to rodent studies designed to demonstrate pharmacologically/genetically mediated alleviations or exacerbations of seizure severity. In studies that employ PTZ induction, seizure severity has often been equated with convulsive severity: *ordinal* scales designed to quantify seizure severity assign low scores to hypoactivity/immobility (assessed empirically), intermediate scores to myoclonic seizures (which are often subtle), and high scores to convulsive or maximal seizures, often with even higher scores for seizures associated with death (15, 16, 19, 22). Our data showed that in the first 30 minutes, PTZ-treated mice (compared with saline) displayed profound immobility and a significant sheltering deficit. In experiments with simultaneous vEEG, PTZ injections in mice have been shown to induce bursts of repetitive (6-7Hz) and/or isolated generalized spike-wave discharges for approximately 15-20 minutes (17, 18). Thus, we hypothesize that this behavioral constellation may represent a transient epileptic encephalopathy, akin to “spike-wave stupor” (57). Since immobility *outside* a shelter (Movie S1) would render a mouse more susceptible to predation, the extent of immobility outside the shelter must be proportional to severity. Interestingly, with subsequent PTZ injections, mice displayed a progressive *improvement* in these early measures (Fig. 2C), albeit at the cost of a progressive increase in the likelihood of convulsion. Convulsive seizures were scored manually: since “spikes” in horizontal displacement or mobility were not always related to convulsions, and since not all convulsions necessarily resulted in such “spikes”, we utilized a trained blinded observer to score for the presence and timing of convulsive seizures.

In the subsequent hour (between 30-90 minutes post-injection), saline-treated mice displayed profound hypoactivity (∼1-2cm/min) and a committed prolonged shelter entry. In contrast, PTZ-treated mice continued to ambulate (∼20-30cm/min), intermittently exited their shelters and entered their food hopper, resembling a “wandering” state (58). This response would *also* render mice vulnerable to predation, and therefore must *also* relate to severity. We observed that repeated PTZ injections did not significantly change these putative measures of wandering (Fig. 2D). Thus, while “convulsive severity” may display a simple linear or step wise worsening daily injections of PTZ (Fig.2F) (16, 17, 19, 20), changes in behavioral severity are more complex. Indeed, both “early” and “late” PTZ-induced seizures were severe in unique ways. Similarly, convulsive and nonconvulsive seizures perturbed home-cage behavior uniquely (Fig. 2G), and aside from the convulsion itself, we found no quantitative evidence to suggest that convulsive seizures are necessarily more “severe”. Distinctions between convulsive and behavioral severity were evident with our other manipulations. In C57BL/6J mice, valproate pretreatment improved convulsive severity without impacting behavioral severity. DBA/2J mice (compared with C57BL/6J) displayed greater convulsive severity (Fig. 5G) while Scn1a^+/-^ mice (compared with wild type littermates) displayed greater behavioral severity (Fig. 6F).

### The Severity of Interictal Behavioral Dysregulation

The second objective of the study was to explore how (i) seizure recency (or frequency) and/or (ii) an increased seizure tendency (i.e., epilepsy) impacts spontaneous home-cage behavior. During the 10-day injection protocol, PTZ-treated mice gradually developed a syndrome of hypoactivity (without excess sleep) and increased overall sheltering behavior. This syndrome was not associated with significant deficits in hedonic drive (sucrose preference, running wheel) or exploratory behavior (light spot, running wheel). Further, this syndrome was transient: it was not evident two months after the cessation of daily PTZ injections. Nevertheless, at this distant time point, persistent differences in “seizure severity” could be demonstrated with a subsequent injection of PTZ. These findings illustrate how interictal behavioral changes that emerge *during* a period of frequent seizures may remit with seizure remission, *without* altering seizure threshold.

Prior to PTZ exposure, DBA/2J mice displayed prominent home-cage hypoactivity compared with C57BL/6J, replicating prior findings (27, 28, 59). In our analysis, hypoactivity could not be explained by excess “sleep” but was associated with (or a result of) fewer but more prolonged feeding bouts. In response to light spot stimulation, DBA/2J mice were markedly different from C57BL/6J mice, with rapid shelter entry and locomotor suppression. While increased sheltering during the light spot test has been advertised as “anxiety-like” (40), mice in the wild naturally *decrease* exploratory behavior under conditions of illumination (e.g., moonlit nights) as an innate response to increased predation risk (36). Thus, enhanced sheltering in response to light spot testing may be a more general measure of *risk aversion*, which may itself be modulated by other factors such as cognitive dysfunction, sympathetic activation and satiety/thirst. Scn1a^+/-^ mice also displayed nocturnal hypoactivity together with a subtle but statistically significant increase in total “sleep”. With a light spot stimulus, suppression of exploration and shelter entry were largely similar between Scn1a^+/-^ mice and littermate controls. In contrast, when mice were transferred (by a human experimenter) into an adjacent cage with novel olfactory cues (“cage swap”), Scn1a^+/-^ mice displayed lower exploratory activity and swifter shelter entry. These data exemplify how differences in exploratory tendency may certainly be test-dependent, and/or may be amplified by human presence.

Nocturnal home-cage hypoactivity that we observed in all three models (repeated PTZ, DBA/2J and Scn1a^+/-^ mice) is itself etiologically nonspecific. Similar observations have been made in some (50–52) but not all (60, 61) mouse autism models, following social defeat stress (62) and also in aged mice (63). Indeed, wide variations in spontaneous home-cage activity and wheel-running exist amongst genetically diverse inbred strains of mice (28, 59). Hypoactivity may either promote survival (by conserving energy and limiting predation) or compromise survival (by lowering access to food/water and mating). Drowsiness and/or encephalopathy could also conceivably result in hypoactivity. However, light spot testing did *not* reveal a blunted or delayed response in either of our three models. Thus, at least during the early active phase (when the light spot was presented), observed hypoactivity was not associated with evidence for an impaired “sensorium” (at least to visual stimuli).

### Limitations

First and foremost, our technique does not (currently) incorporate simultaneous electroencephalography (EEG). Thus, our approach fails to provide a temporally precise appreciation of how the presence or absence of intermittent epileptiform discharges may correlate with putative “stuporous” or “wandering” behavior. In the same light, we are unable to discern how the timing of spontaneous seizures in adult Scn1a^+/-^ mice (albeit rare (49)) may transiently alter home-cage behavior. Combining long term home-cage monitoring with synchronized high-quality wireless EEG remains an important ongoing objective but may reveal *observer effects*: the mere placement of EEG electrodes in mice (tethered *or* wireless) may impair mobility, sleep and neurovegetative function through pain or direct physical hindrance. Further, EEG implantation surgery may itself increase seizure threshold through pain, impaired sleep and postoperative inflammation (64), as well as through unintended cortical microlesions that may occur when electrodes are inadvertently advanced to subdural depths (65).

Second, this report describes seizures artificially induced by a chemoconvulsant. We chose this approach, as it allowed us to induce seizures “on-demand” in large cohorts of mice, thereby synchronizing our assessment of ictal and interictal behavior. It also enabled us to model a brief period of frequent seizures followed by remission. While acute seizure protocols are invaluable to screen and identify novel anticonvulsants (12), their relevance to infer the pathophysiology of epilepsy is admittedly inferior to models that display frank spontaneous seizures. Since PTZ is a GABA-A receptor antagonist (13), differences in seizure severity may be a reflection of primary alterations in GABA-A receptor expression and/or subunit composition. These differences may be a critical component of the enduring seizure predisposition in some but not all models of epilepsy. Future home-cage experiments that incorporate pharmacologically distinct chemoconvulsants or other clinically relevant seizure triggers (e.g., sleep deprivation or fever), may offer additional insights that may not be evident with PTZ.

### Conclusions

As we continue to expand our knowledge of epilepsy genetics, innovate mouse models that recapitulate those genetic lesions, and apply novel anatomically targeted methods to interrogate network dysfunction in epilepsy, we must continue to refine our assessment of seizure severity and interictal behavior in preclinical models. Our data illustrates the utility of continuous long-term home-cage monitoring in assessing these important components of epilepsy disability in mice. Future studies that incorporate home-cage monitoring may help dissect how interictal behavior is modulated by seizure onset (focal Vs generalized), spontaneous seizure or spike frequency, as well as discern why certain anticonvulsants, regardless of epilepsy type or seizure frequency, are themselves associated with high rates of adverse behavioral side effects (66, 67).

## Materials and Methods

### Mice and Drugs

This study was conducted in accordance with the Institutional Animal Care and Use Committee at Baylor College of Medicine. Mice were housed with a 12-hour light cycle (lights ON from 0500-1700) under controlled temperature (20-26°C) and humidity (40-70%) conditions. Pups were weaned at ∼3 weeks of age and group-housed by sex. Mice were provided *ad libitum* access to water, chow (Pico Lab ® Select Rodent 5V5R Standard Diet) and were housed on Biofresh Performance Pelleted Cellulose bedding. C57BL/6J and DBA/2J breeders (#000664 and #000671, Jackson Laboratories) obtained in October 2017 were used to generate respective colonies. Scn1a R1407X mice (47) were genotyped by polymerase chain reactions of tail DNA (46, 47): wildtype-forward (5’-ATGATTCCTAGGGGGATGTC-3’), mutant-forward (5’-TTTACTTTCACATTTTTCCATCA-3’), and common reverse (5’-CTTTCACATTTTTCCACCG-3’). All drugs were injected intraperitoneally (5ml/kg). Pentylenetetrazole and sodium valproate (Sigma-Aldrich) were dissolved in sterile-filtered normal saline for a final dose of 30-60mg/kg and 200mg/kg respectively. Diazepam (3mg/kg, Henry Schein) was dissolved in manufacturer-provided diluent (propylene glycol) (68).

### Home-cage Monitoring

At the start of the experiment, mice were transferred into a satellite facility with lighting, humidity and temperature settings identical to vivarium conditions. A sound machine (‘Lectrofan, Adaptive Sound Technologies) played continuous white noise. Satellite access was restricted to experimenters (VK, MJ) who wore personal protective equipment (gown, cap, face mask and gloves) and entered once daily to visually inspect food and water sources. Veterinary inspections were performed every other week (∼1100-1200). Mice were individually housed in one of sixteen Phenotyper ® (Noldus Information Technology) home-cages designed for mice (30×30×47cm) with clear plastic walls customized to incorporate two lickometered water sources (detecting capacitance changes), a feeding meter (detecting beam breaks) and a detachable running wheel (utilizing a wheel-attached magnet and a wall-attached magnet sensor). Chow and drinking water were identical to those provided in vivarium cages, and sucrose (Sigma-Aldrich) was dissolved directly into fresh drinking water at a final concentration of 0.8% (w/v). An infrared-lucent shelter (10×10×5-6cm) was also employed.

### Videotracking, Recordings and Injections

An aerial video-feed was established using a ceiling-mounted infrared camera together with an array of ceiling mounted infrared lamps. A “shelter zone” was defined on a distance-calibrated “arena” for each cage. Live video-tracking was conducted via Ethovision XT 14 (Noldus) using dynamic subtraction at a sample rate of 15/s. Licking, feeding and wheel-running data were integrated through a hardware control module. Detection settings were consistent for all mice of any given experiment. Recordings of home cage behavior were initiated either as a “manual” start (i.e., start “now”, for ictal recordings) or started at a specific “clock” time (for baseline or interictal recordings). The “light spot” was produced by a white spotlight resulting in fairly diffuse cage illumination centered in the upper left quadrant of the cage. For the “cage-swap” experiment (Fig. 6E), mice were “swapped” into a cage previously inhabited by a sex-matched conspecific for a period of 2h and then swapped back into their original cages. To validate our “sleep” measurement algorithm, a separate group of C57BL/6J mice were acclimated to home-cages for 48h and then received a single injection of 3mg/kg diazepam just prior to 1700h (Fig. S1A).

### Data Acquisition

For each epoch of recording (∼2-3h or ∼20h in duration), data pertaining to horizontal distances (resulting in a displacement of *x* and/or *y* coordinates) was automatically tallied by Ethovision XT 14 and configured to ignore horizontal displacements of less than 0.2cm (between samples). Such “track-smoothing” was necessary to avoid out oscillatory tracking “noise” in mice that were either extremely immobile or deceased. Distance, licking duration, feeding duration (or entries) and sheltering time were reported as daily totals (for time budgets) or binned as either total/minute or total/hour. Convulsive seizures were defined as any abrupt paroxysm that resulted in a loss of upright posture, clonic and/or tonic movements, with or without wild running, and/or hind limb extension (Movie S3, S4). Convulsive events were identified by screening “ictal recordings” through integrated visualization (Movie S2), where changes in “% mobility” (defined as the percentage of *changed pixels* of an object between samples) were reviewed simultaneously with video data. For each “ictal” recording, all spikes in % mobility were screened and the latencies of true convulsive seizures (confirmed by video review) were reported.

### Data Analysis

Graphs and statistical analyses were generated on Prism GraphPad 8. To assess total sleep time and sleep bout duration/occurrence, a stand-alone module was designed to analyze Ethovision raw data files and identify 40s-long contiguous periods devoid of “movement” (defined as sample velocity ≤1.2cm/s). When mice died following a PTZ injection, death was tallied in survival curves (Fig. 2F, Fig. S5). However, data from that specific PTZ injection was excluded from any graphs of averaged “ictal” behavior (e.g., Fig. 2A, 2G or 5G). These deaths also resulted in missing values for daily total measurements (e.g., Fig. 2B or 3B). To perform repeated measures analysis of variance (RMANOVA) with missing values, a mixed effects model was employed, and interactions between fixed effects are reported (e.g., *group x time*, or *group x day*). Unpaired two-tailed Student’s *t* tests were used to compare two group means (e.g., Fig. 2E or 4E). Two-tailed Chi squared analyses were employed to demonstrate significant changes in the *fractions* of mice that convulsed.

**Movie S1:** An example of early immobility outside the shelter. In this video (which captures behavior ∼7.5 minutes post-injection), the mouse on the top right chamber displays a bout of prolonged immobility following PTZ. Saline-treated mice (all others in this video) are seen within their shelters either grooming or resting.

**Movie S2:** Integrated visualization for the detection of convulsive seizures. In this video, aerial footage from home-cage chambers are shown on the left with simultaneous measures of “% mobility” on the right (defined as the percentage of changed in pixels of an object between individual samples). In this video, the mouse in chamber PT14 (top right) experiences a convulsive seizure with hindlimb extension marked by a polyphasic transient change in mobility.

**Movie S3:** An example of a convulsive seizure outside the shelter (top right chamber).

**Movie S4:** An example of a convulsive seizure within the shelter (bottom right chamber).

## Acknowledgments

We thank Drs. Jeffrey Noebels, Atul Maheshwari and Joachen Meyer for their editorial and scientific comments

## Author contributions

V.K. conceptualized the experiments and wrote the paper. M.J.J and V.K. performed the experiments. P.P.K. designed and implemented a module to detect and quantify sleep in mice. Scn1a^+/-^ and WT littermate breeding pairs (47) were generously provided by Dr. Jeffrey Noebels (46).

## Data and materials availability

All data associated with this paper are in the main text or the supplementary materials.

## Supplementary Materials

**Fig. S1.**
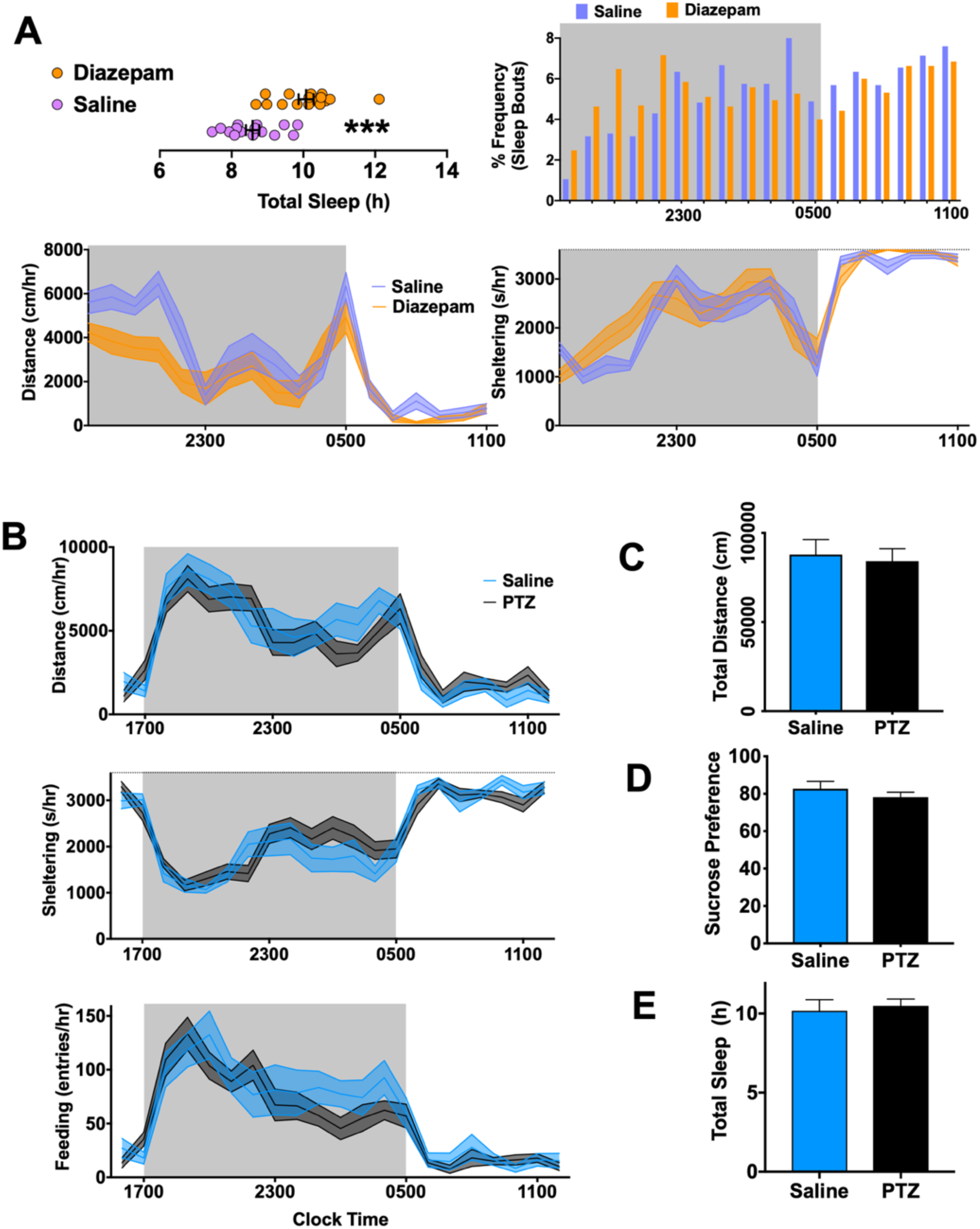
**(A)** A separate group of 8-10week old C57BL/6J mice were acclimated to home cage chambers for two days and then received either an injection of saline (n=15, 7F) or diazepam (3mg/kg, n=16, 8F) at ∼1655. Diazepam-treated mice displayed significantly greater average total sleep with a prominent increase in sleep bouts between 1700-2300. Diazepam also reduced overall distances moved (group x hour, F_17,493_= 2.1, p<0.01) and increased sheltering (group x hour, F_17,522_=1.7, p<0.05). **(B)** After two consecutive days of baseline recording, C57BL/6J mice (from Fig. 1) were randomized to receive saline (n=12) and PTZ (n=20). Distances (group x hour, F_20,600_=1.3, p>0.1), sheltering (group x hour, F_20,600_=1.3, p>0.1) and feeding entries (group x hour, F_20,600_=1.1, p>0.1) were not significantly different. **(C-E)** Groups were also similar in total distance moved, sucrose preference and total sleep. ***: p<0.01. Mean ± SEM shown.

**Fig. S2.**
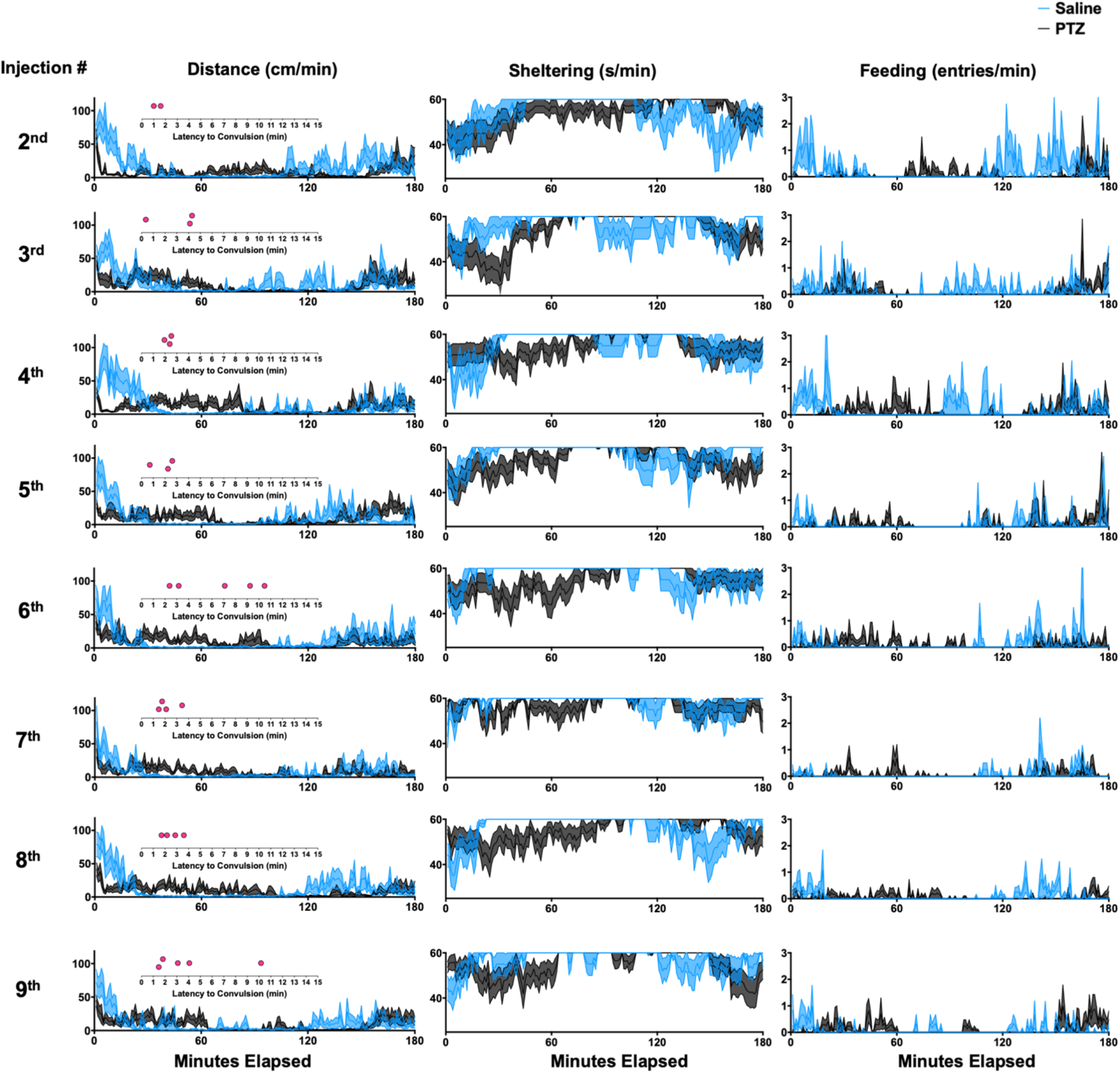
Injections of saline Vs PTZ (30mg/kg). LEFT (distances), CENTER (sheltering) and RIGHT (feeding entries) for the 2^nd^ through 9^th^ injections with dot plots reflecting the latency to convulsions in each ictal recording. Mean ± SEM shown.

**Fig. S3.**
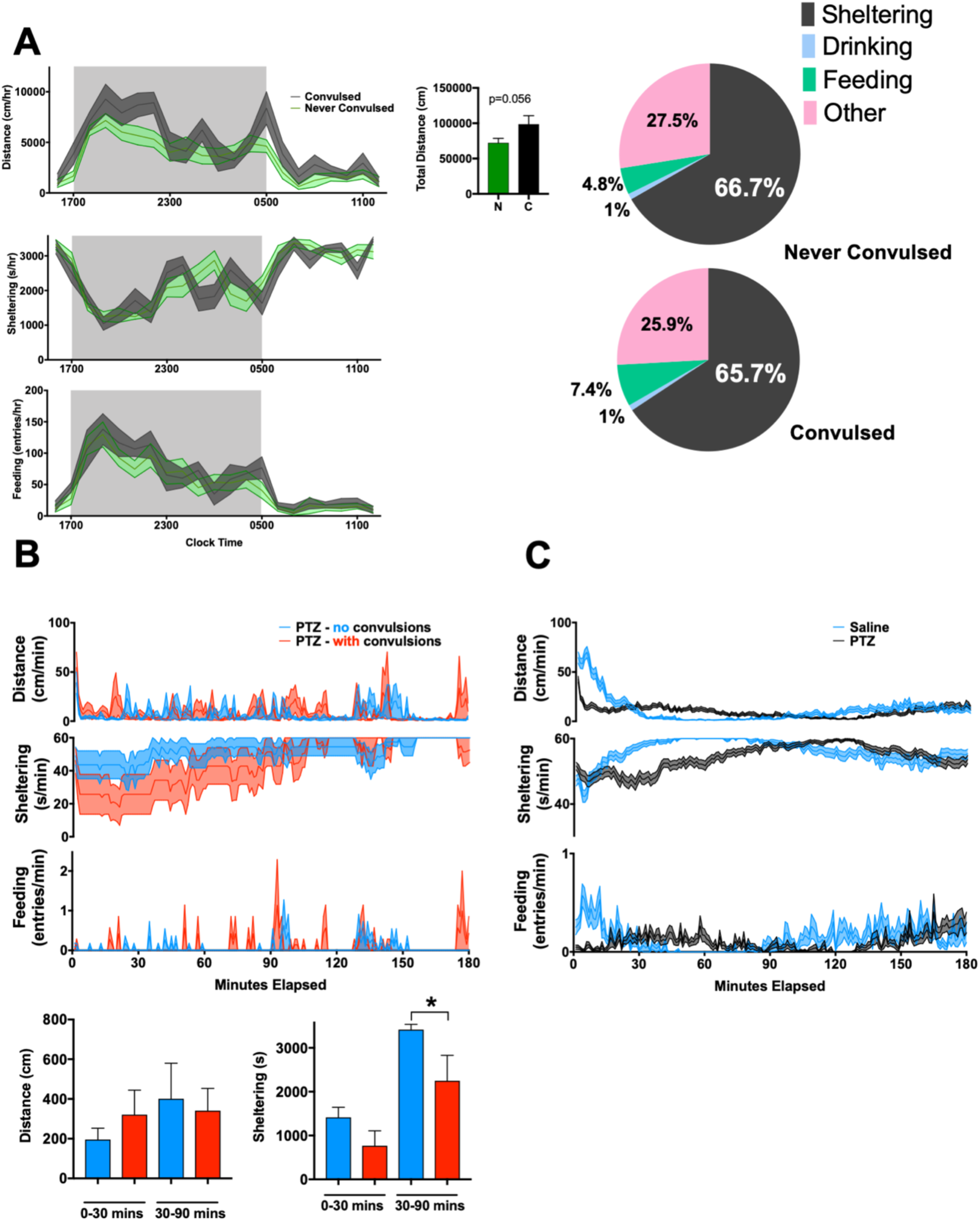
**(A)** Over the 10-dayprotocol, 9 mice displayed convulsions following any PTZ injections. On their second baseline day (Fig. 1), compared with the 11 mice that never convulsed, these mice displayed a trend to accumulate greater distances (group x hour F_20,360_=1.4, p=0.1), altered sheltering patterns (group x hour F_20,360_=2.0, p<0.01) without changes in feeding entries (group x hour, F_20,360_=0.5, p>0.9). RIGHT: Time budgets for these groups. **(B)** Behavioral response to the FIRST injection of PTZ in mice that never convulsed (n=11) and those that ever convulsed (n=7). We exclude two mice that convulsed following the first injection. **(C)** Averaged across all ten days, PTZ injections produced a mean behavioral response that was clearly distinct from saline. *: p<0.05. Mean ± SEM shown.

**Fig. S4.**
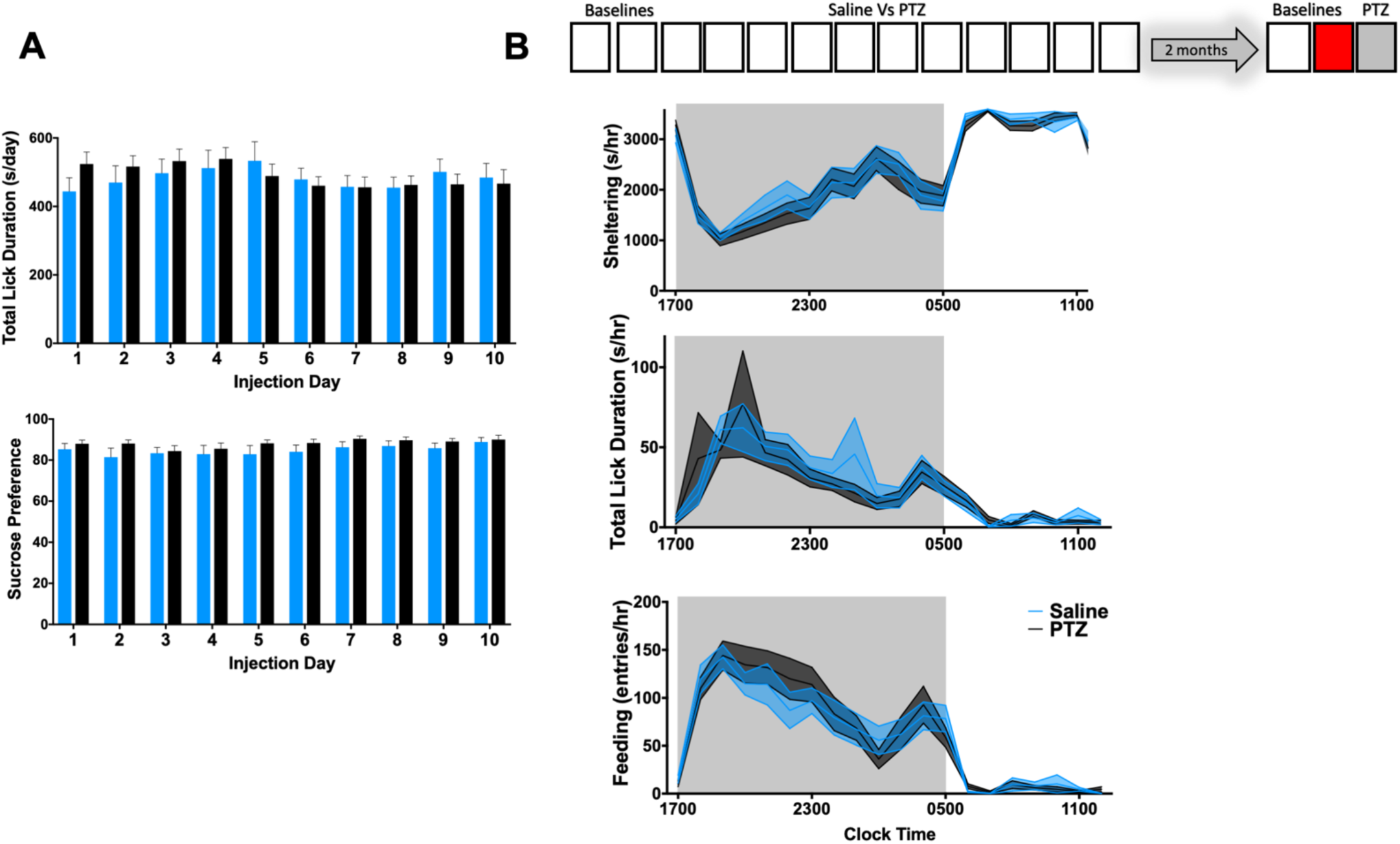
**(A)** Across 10 “interictal” recordings (1600-1100, corresponding to data in Fig. 3B), PTZ- and saline-treated mice displayed similar total licking (group x day, F_9,261_=1.1, p>0.1) or sucrose preference (group x day, F_9,261_=0.4, p>0.5). **(B)** Other behavioral parameters measured (for Fig. 4B) showing no differences between saline and PTZ-treated mice on measures of sheltering (group x hour, F_19,513_=0.3, p>0.9), licking (group x hour, F_19,513_=0.4, p>0.9) or feeding (group x hour, F_19,513_=0.5, p>0.9). Mean ± SEM shown.

**Fig. S5.**
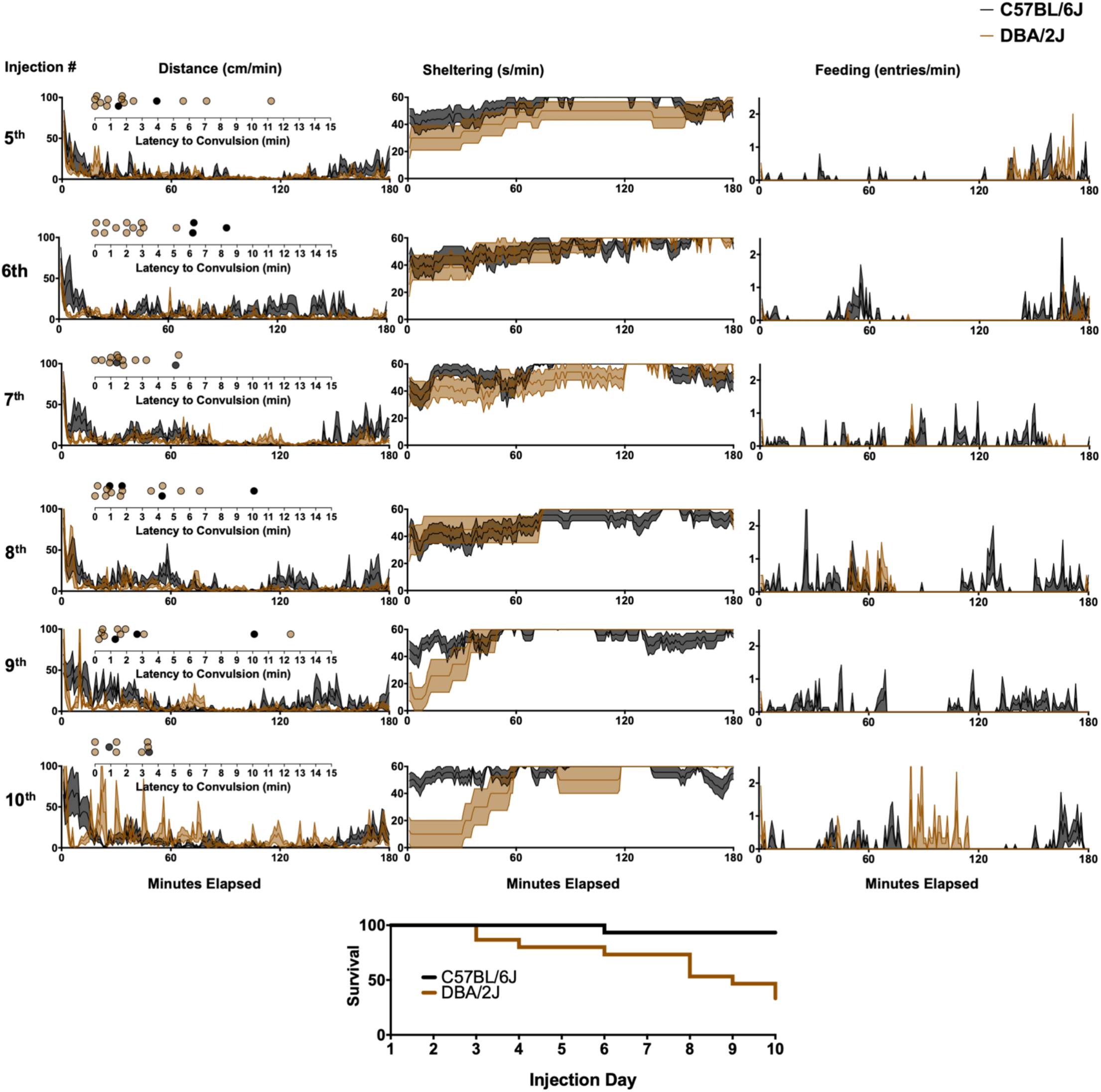
Injections of PTZ in C57BL/6J Vs DBA/2J. LEFT (distances), CENTER (sheltering) and RIGHT (feeding entries) for the 5^nd^ through 10^th^ injection with dot plots reflecting the latency to convulsions in each ictal recording. BOTTOM: Overall survival curve (includes ictal recordings depicted in Fig. 5). Mean ± SEM shown.

